# Light-orchestrated multi-step solid-phase picodroplet reactors

**DOI:** 10.1101/2025.06.23.660884

**Authors:** Mo Wu, Mohammad A. Zaman, Michael A. Jensen, Punnag Padhy, Wei Ren, Ronald W. Davis, Lambertus Hesselink

## Abstract

Picoliter droplet reactors, enabled by droplet microfluidics, are revolutionary tools for biochemical reactions with high efficiency, precision, and minimal reagent use. They excel in single-step reactions and reagent addition through droplet merging. Integrating reagent exchange and washing capabilities directly into these platforms may enable complex multi-step processes, such as de novo oligonucleotide synthesis and multiplex immunoassays. However, current picodroplet microfluidics, lacking such capabilities, remain critically deficient in executing multi-step processes. Here we introduce light-orchestrated solid-phase picodroplet reactors to overcome these limitations. Our platform employs optoelectronic tweezers to manipulate individual picodroplets and microbeads. Using this platform, we demonstrate an eight-step click chemistry-based DNA ligation synthesis cycle, with real-time in situ fluorescence detection of reaction products. This platform overcomes single-step limitations by achieving precise sequential encapsulation and decapsulation of beads with different reagent droplets, ensuring uniform reagent exposure and effective washing. Hence it mitigates reaction errors caused by nonuniform reagent exposure and trapped impurities in conventional bulk processing of microbeads. With minimal reagent consumption, real-time analysis, and programmable light-based control, the platform can potentially be scaled and fully automated into a universal and versatile picoliter-scale reagent handling robot for miniaturizing and streamlining workflows in synthetic biology, drug discovery, and beyond.

## Introduction

Picodroplet reactors – platforms for performing picoliter-volume reactions enabled by droplet microfluidics – are emerging as powerful tools for biochemical reactions with minimal reagent use and high precision^1–5^. With established droplet processing techniques including merging, splitting, mixing, incubation, and sorting, picodroplet reactors are versatile for biochemical workflows such as single-cell analysis and high-throughput screening for drug discovery and directed evolution^6–8^. However, many other important biochemical workflows, such as oligonucleotide synthesis, enzymatic assays, and immunoassays, require multi-step processes involving multiple reagent exchanges, which entail the sequential addition and subsequent removal of different reagents, as well as product extraction. Current picodroplet platforms cannot support such multi-step workflows, as they lack reliable mechanisms for controlled reagent exchange^3^.

Typically, multi-step biochemical workflows are performed on solid-phase platforms in column or multi-well plate formats^9–11^. In such platforms, microbeads (e.g., polystyrene or glass) are commonly used as solid supports to provide surfaces where reactant molecules can bind and undergo biochemical reactions. Because the reaction products remain tethered to the bead surface, product extraction is simplified, enabling high-throughput processing^12,13^. Despite these advantages, conventional solid-phase platforms face significant limitations. Processing millions of beads collectively in bulk leads to nonuniform reagent exposure and trapping of residual impurities within the bead matrix, resulting in incomplete reactions and error propagation across multiple reaction steps and cycles^14–16^ (Fig. 1, a).

**Fig. 1:**
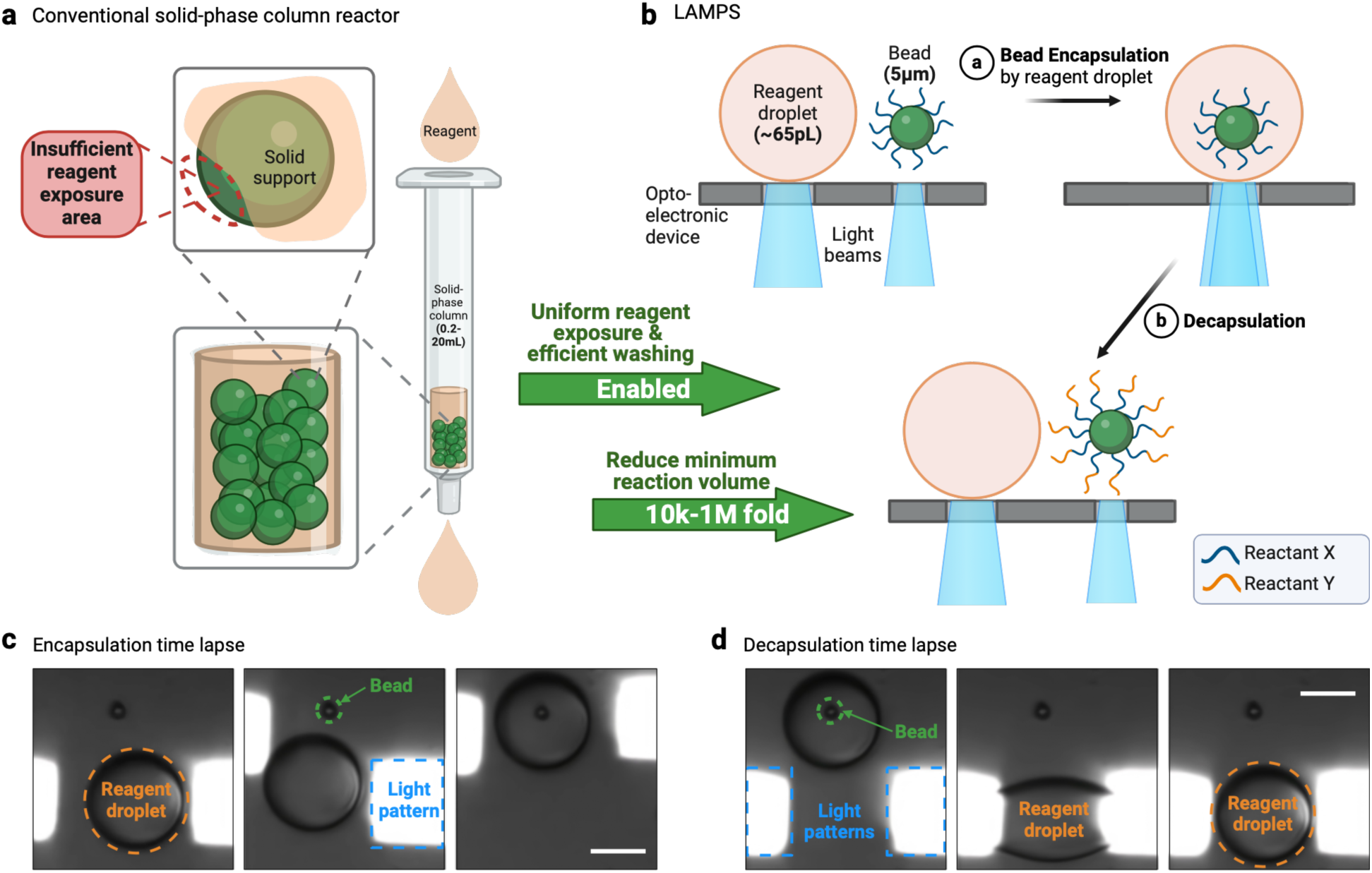
Encapsulation and decapsulation mechanism for reaction on LAMPS and its advantages. **a**, Limitations of nonuniform reagent exposure on conventional platform. Reaction errors can be introduced by insufficient reagent exposure area on the solid supports (i.e., microbeads) and impurities trapped within the bead matrix. **b**, Main feature of LAMPS: single bead is encapsulated and decapsulated by reagent picodroplet. The reaction is initiated when the bead (coated with reactant X) is encapsulated by the corresponding reagent droplet (containing reactant Y) and terminated when the bead is decapsulated by the droplet. The bead is 5 µm in diameter and the droplet is ∼65 pL in volume (50 µm in diameter). **c, d**, Experimental images of encapsulation and decapsulation of a microbead by a reagent picodroplet. For decapsulation, the droplet is pulled towards the light patterns during the active period of the voltage, and retracts to its spherical shape during the inactive period. Scale bars: 25 µm.

While not originally developed to address limitations of conventional solid-phase systems, picodroplet microfluidics has been explored to encapsulate and isolate individual reaction sites (i.e., microbeads) within droplets. These efforts have focused primarily on enhancing detection sensitivity, enabling single-molecule assays, and performing digital quantification in biochemical and diagnostic applications through compartmentalization^17,18^. Although these systems can achieve significant sensitivity gains by reducing reaction volume and enhancing signal-to-noise ratios^19,20^, they again remain limited to single-step reactions due to the lack of mechanisms for reagent removal or bead decapsulation. Without the ability to exchange reagents between reaction steps, multi-step biochemical workflows remain inaccessible in these picodroplet systems.

To overcome these limitations and enable multi-step solid-phase reactions in picodroplet systems, we developed a platform of light-actuated multi-step picodroplet solid-phase reactors (LAMPS). The LAMPS platform leverages optoelectronic tweezers (OET) to achieve programmable, sequential encapsulation and decapsulation of individual microbeads by corresponding reagent picodroplets (Fig. 1, b to d). It is a microfluidic optoelectronic device that integrates on-demand droplet generation with nondestructive controlled manipulation of picodroplets and beads. Individual picoreactor droplets are trapped and manipulated in parallel using programmable micro light patterns that are projected onto the device. It enables the reaction exchange capabilities that are missing in current droplet microfluidics. The LAMPS platform has several advantages over conventional solid-phase reactors as well. It ensures uniform reagent exposure, enhances reaction efficiency, and facilitates more effective washing steps between reaction steps. Additionally, replacing microtiter plates with picodroplet reactors allows reactions to be performed at significantly smaller volumes. The minimum reaction size is reduced from microliters to picoliters, minimizing reagent use and waste by up to a million-fold. By integrating reagent exchange capabilities into picoliter droplet microfluidics, the LAMPS platform enables solid-phase reactions in picodroplet reactors without inheriting the limitations of conventional solid-phase reactors.

We designed LAMPS to be a picodroplet microfluidic platform, which can host a variety of multi-step solid-phase reactions. To demonstrate its capabilities, we performed a modified DNA ligation synthesis cycle based on copper-catalyzed azide-alkyne cycloaddition (CuAAC) click chemistry. The reaction cycle consists of an azide addition reaction step using terminal deoxynucleotidyl transferase (TdT), followed by a CuAAC click ligation reaction step, with three washing steps performed after each reaction step. The washing steps are essential for removing spent reactants from the previous step. This eight-step ligation cycle enables precise stepwise assembly of sequence-verified oligonucleotides on the LAMPS platform, demonstrating its ability to handle diverse reagents and execute multiple sequential reaction steps on individual beads.

## Results

### LAMPS platform design

The LAMPS platform, engineered as a solid-phase picodroplet reactor, enables reliable encapsulation and decapsulation of microbeads with reagent droplets. This ensures uniform reagent exposure over the surface of each bead followed by complete reagent removal postreaction. The LAMPS device consists of an optoelectronic trapping/manipulation surface bonded to a polydimethylsiloxane (PDMS) layer hosting the microfluidic components. A central reaction chamber (∼ 5 mm × 5 mm) along with oil (i.e., suspension medium) inlet/outlet and four on-demand droplet generators are patterned in the PDMS layer. In our setup (Fig. 2a), 5 µm diameter magnetic beads, acting as solid supports, are suspended in the oil medium of hexadecane. This colloidal solution is then introduced into the central chamber of the device.

**Fig. 2:**
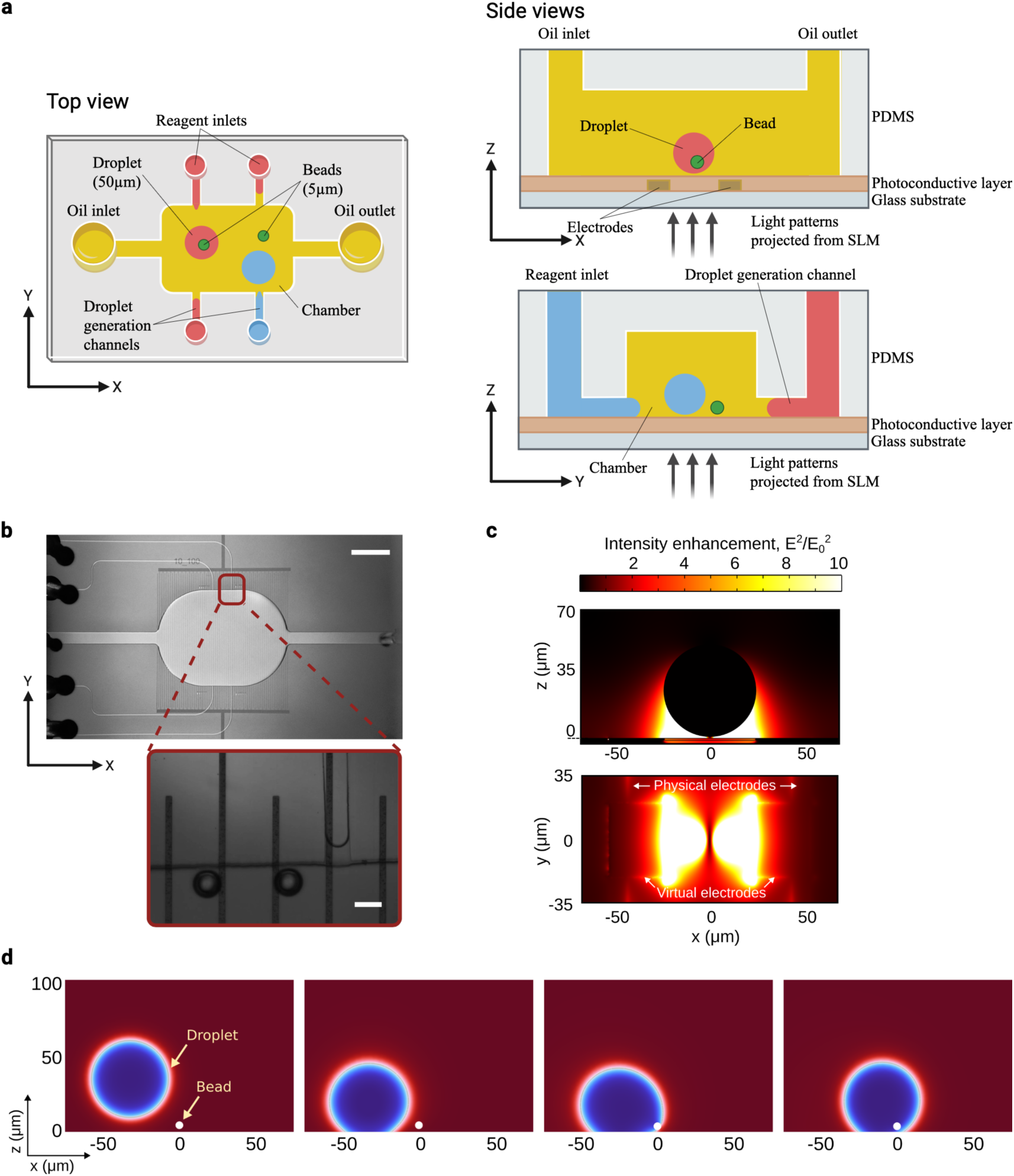
LAMPS platform design. **a**, Schematic showing top and side views of the LAMPS platform. Hexadecane is used as the oil medium in this study. **b**, Top-view optical image of the device. Four microfluidic droplet generation channels, each for a different reagent, are connected to a central chamber. Reagent droplets are generated at the intersections of the microfluidic channels and the central chamber (inset). Scale bars: 2 mm; 50 µm (inset). **c**, Numerical simulation of the electric field. **d**, Numerical simulation of the droplet motion during bead encapsulation. The gradient force exerted on the droplet from the light-activated virtual electrodes is sufficient to overcome its surface tension.

Hexadecane is chemically inert, optically transparent, and immiscible with the reagent chemicals involved in the CuAAC-based DNA ligation synthesis cycle, making it an ideal suspension medium. We have experimentally confirmed its compatibility with aqueous solutions and prevalent solvents used in biochemical reactions, such as ethanol, dimethylformamide (DMF), and tert-butanol. Droplets containing ∼65 pL of reagent (50 µm in diameter) are generated on-demand into the chamber through microfluidic channels driven by a piezoelectric pressure controller. The step-emulsification droplet generation mechanism employed here depends primarily on the channel geometry and is not overly sensitive to pressure fluctuations. This ensures consistent on-demand droplet generation when using a standard benchtop pressure controller^21–23^. The full experimental setup schematic is shown in Extended Data Fig. 1.

The parallel trapping and manipulation of the picoreactor droplets is carried out using adapted optoelectronic tweezers (OET), a technology known for its flexibility and versatility in microparticle manipulation^24–26^. An electric field gradient, created by light-activated virtual electrodes on a photoconductor, exerts trapping force on dielectric particles and droplets that have different material properties (i.e., permittivity and electrical conductivity) compared to the suspension medium. The low optical power requirement of OET makes them especially suitable for handing sensitive biomolecules that are prone to photothermal damage. Additionally, the low electrical conductivity of the suspension medium hexadecane mitigates the Joule heating effect that can also be detrimental to the biomolecules. On the other hand, the low conductivity of hexadecane makes the conventional vertical electrode configuration of OET ineffective for creating electric field gradients^24,27^. Thus, we adapted an OET device with a planar electrode layout. Our OET device features a 1 μm thick photoconductive layer of hydrogenated amorphous silicon (a:Si-H) deposited over gold interdigitated microelectrodes. This forms the optoelectronic trapping/manipulation surface. The interdigitated electrodes, with a width of 10 µm and spacings of 100 µm, are designed to work optimally for the 50 µm diameter picodroplets (Fig. 2b). Micro-sized light patterns are projected onto the photoconductive layer of the device by coupling the output of a spatial light modulator (SLM) to a custom inverted microscope, thereby creating localized high conductivity regions within the photoconductive layer. These regions, referred to as “virtual electrodes”, behave akin to the physical microelectrodes used in dielectrophoresis systems. With an AC voltage (9.3 V, 10 kHz) applied across the interdigitated electrodes, a nonuniform electric field is created near the virtual electrodes. The resulting field gradient is significant on the same spatial scale as the beads and the droplets and, therefore, can exert sufficient trapping forces on them. This is modeled through numerical simulations of the electric field (Fig. 2c) and the droplet motion (Fig. 2d) under the influence of the OET gradient forces (Extended Data Figs. 5 and 6), as well as verified from experimental observations (Fig. 1, c and d).

Employing a programmable SLM to steer the micro light patterns in real time, the LAMPS platform is capable of dynamic control, both spatial and temporal, of the trapping force, achieving precise and automated two-dimensional parallel manipulation of picodroplets (Fig. 3). Fig. 1c and Fig. 1d are images taken during experiment highlighting the moments of bead encapsulation and decapsulation, respectively. Once a droplet is generated near the microfluidic channel opening, two rectangular micro light patterns are used to seize the opposite ends of the droplet and subsequently guide its trajectory towards the trapped bead by translating the light patterns. Precise control over the light pattern translation is achieved through custom software to control the SLM. Desired light-pattern trajectories are preprogrammed to achieve semi-automated motion. As the droplet is pushed onto the bead surface, the applied gradient force is sufficient to overcome the surface tension of the droplet, leading to complete encapsulation of the bead (Figs. 1c, 2d, and 3a). This ensures that the bead is fully exposed to the reagent over its entire surface during the reaction period. In addition, moving the droplet with the encapsulated bead induces fluid flow inside the droplet^28,29^; this simulates a stirring mechanism that promotes reagent mixing for consistent bead surface exposure, which helps drive the reaction to completion.

**Fig. 3:**
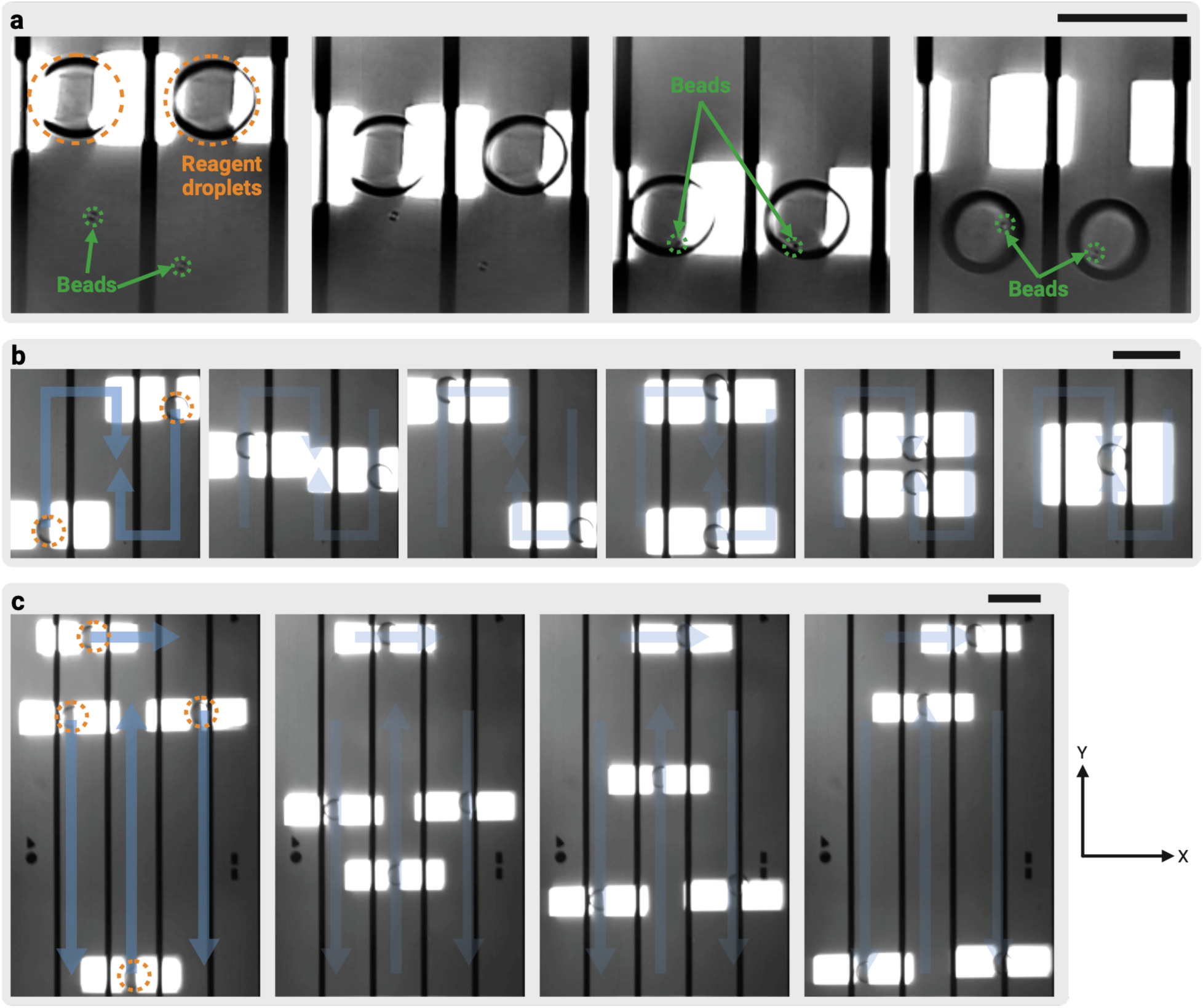
Automated parallel 2D manipulation of picoreactor droplets achieved by LAMPS. **a**, Parallel encapsulation of two microbeads. **b**, Parallel manipulation and merging of two picodroplets. The two droplets move along preprogrammed paths that converge, resulting in their merging into a single droplet. **c**, Simultaneous parallel manipulation of four picodroplets in different directions along the preprogrammed paths. The preprogrammed paths are delineated by light blue arrows, and the droplets are outlined with orange dashed circles in (**b**) and (**c**). Scale bars: 100 µm.

To initiate decapsulation, the pair of rectangular light patterns are placed slightly away from the droplet and a pulse-modulated sinusoidal voltage (duty cycle 50%, period 2 s) is applied (Fig. 1d). This tailored voltage pattern triggers pronounced stretching and contraction of the droplet between the light patterns, a phenomenon attributable to both dielectrophoresis and electrowetting effects^30–32^. The stretching reduces the localized curvature of the droplet and enables bead decapsulation. During the period of reduced voltage, the droplet seeks to revert to its spherical shape due to surface tension, a characteristic that helps maintain droplet integrity and aids the decapsulation procedure.

### Multi-step oligo assembly demonstration

By combining CuAAC click ligation and enzymatic azide addition, we have developed an eight-step DNA ligation synthesis cycle on the LAMPS platform (including washing steps), depicted in Fig. 4a. The cycle is designed to be iterated to concatenate sequence-verified oligonucleotides in a controlled manner for de novo DNA synthesis. In situ fluorescence analysis confirms the successful execution of the full reaction cycle (Fig. 4, b and c; see Extended Data Fig. 2 and Extended Data Fig. 3 for detailed fluorescence analysis).

**Fig. 4:**
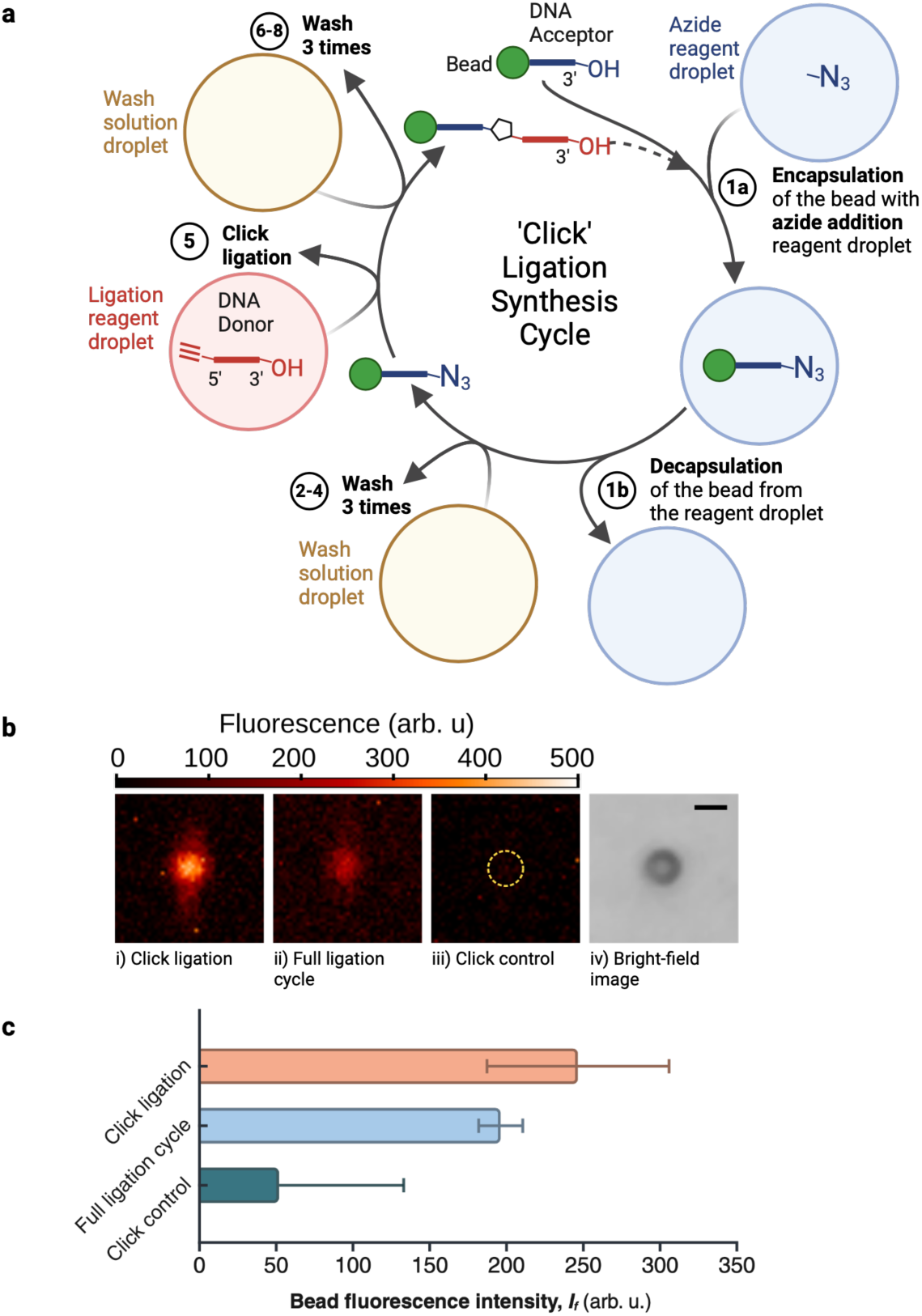
LAMPS demonstration with click ligation synthesis cycle. **a**, Eight-step template- independent DNA click ligation synthesis cycle on the LAMPS platform. Each step is performed by encapsulation and decapsulation of the microbead with the corresponding reagent picodroplet. Step 1: Azide addition. 3’-azide modified dNTP is coupled to DNA acceptor molecule using TdT. Steps 2 to 4: Washing steps with three PBS droplets. Step 5: Click ligation. 5’-alkyne modified donor molecule is ligated with 3’-azide acceptor molecule through CuAAC click chemistry. The ligated product has a triazole linkage. Steps 6 to 8: Washing steps with three PBS droplets. **b**, Fluorescence images of beads after three different reactions: (i) Click ligation step only with 3’-azide acceptor molecule; (ii) full ligation cycle, including both azide addition and click ligation steps, with 3’-hydroxyl acceptor molecule; and (iii) a negative control with 3’- hydroxyl acceptor molecule, where only the click ligation step was performed without the azide addition step and no ligation is expected. Low to no fluorescence in the control sample establishes a baseline for comparison. The fluorescence signal emitted by the ligated donor molecule labeled with Atto 647 after click ligation or full ligation cycle confirms successful reactions. (iv) A bright-field image of a representative bead is shown for reference. Scale bar: 5 µm. **c**, Bar plot of the mean fluorescence intensity of bead samples from LAMPS after the three reactions. The error bars show the standard deviations.

### CuAAC click chemistry-based DNA ligation

Traditional enzymatic DNA ligation methods face limitations such as the requirement for a template, difficulty joining fragments with repetitive sequences, and reduced compatibility with modified nucleotides^33,34^. Chemical ligation methods, however, may overcome these challenges and offer flexibility for diverse artificial modifications to DNA molecules^35,36^. The CuAAC click reaction facilitates the creation of covalent bonds between azide and alkyne groups. This enables the seamless integration of various molecular entities and the synthesis of complex molecules that are difficult or impossible to achieve using traditional methods^37^. Click DNA ligation with triazole formation has emerged as an important technique for synthesizing long DNA strands^33,38^. Although the triazole linkage is different from the natural phosphate backbone of DNA, it is compatible with PCR and various downstream applications^38–40^. To illustrate the capability of LAMPS for template-independent DNA assembly, we perform solid-phase CuAAC click ligation in picodroplets (Fig. 4a, Step 5).

Magnetic beads (diameter of 5 µm) as solid supports are initially coated with ∼500 attomoles of 3’-azide-modified DNA acceptor molecule. The donor molecule is a sequence-verified oligonucleotide modified with a 5’-alkyne and an internal Atto 647 fluorescent label. The 3’-end of the donor molecule is unmodified, retaining a hydroxyl group to prevent self-ligation. To avoid side reactions involving the 5’-alkyne of the donor molecule, copper(I) catalyst, and oxygen^41,42^, a droplet containing the donor molecule and another droplet containing the copper(I) catalyst are generated on-demand and introduced separately. The two droplets are then brought together with light patterns to merge and mix on-chip, forming a mixture droplet with minimized oxygen exposure. A single bead is encapsulated by the mixture droplet for the duration of the click reaction. The reaction is then terminated by decapsulation of the bead, followed by three sequential washing steps in which fresh phosphate-buffered saline (PBS) droplets are introduced in succession to encapsulate and decapsulate the bead; this removes any non-specific binding of unreacted donor molecule (Fig. 4a, Steps 6 to 8). The success of the click ligation reaction is verified in real time by in situ measurement of the fluorescence signal from the bead, which is emitted by the ligated donor molecule labeled with Atto 647 (Fig. 4, b and c).

### TdT-based enzymatic azide addition

After the click ligation step, the DNA molecule bound to the beads have a 3’-hydroxyl group, which blocks further click reactions. To enable the addition of subsequent strands through click ligation, an azide addition step is incorporated into the reaction cycle (Fig. 4a, Step 1). This step is performed by the enzymatic action of TdT, a template-independent DNA polymerase that adds nucleotides to the 3’-end of a DNA strand^43,44^. With the 3’-azide modified deoxynucleotide triphosphate (dNTP), TdT couples precisely one nucleotide to the acceptor molecule, as the 3’- azide blocks additional nucleotide coupling and prevents uncontrolled polymerization. The completion of the enzymatic coupling reaction is then verified on the LAMPS platform, where the bead is encapsulated and decapsulated by an enzymatic addition droplet, followed by three sequential PBS washing steps to remove unreacted components (Fig. 4a, Steps 1 to 4). Moreover, the physical properties of the click ligation droplet and the azide addition droplet differ significantly. The ability to orchestrate both kinds of droplets demonstrates the robustness and versatility of the LAMPS platform.

### Click ligation synthesis cycle

By combining enzymatic azide addition with subsequent CuAAC click ligation, the LAMPS platform enables a template-independent, stepwise DNA ligation synthesis cycle to concatenate sequence-verified oligonucleotides within picodroplets. To demonstrate a full ligation cycle on the LAMPS platform, the acceptor molecule initially bound to the beads is unmodified with a 3’- hydroxyl group, the dNTP used in the enzymatic azide addition step has a 3’-azide group without fluorescence labeling, and the donor molecule is modified with a 5’-alkyne group and internal Atto 647 label. Sufficient washing after each reaction step, performed by three sequential fresh PBS droplets, was found to be essential for completing the full reaction cycle. The success of the ligation cycle is verified by in situ detection of a fluorescence signal from the Atto 647-labeled donor molecule (Fig. 4, b and c). In this case, a fluorescence signal is only possible if azide was coupled to the acceptor, which makes it an accurate indicator of the reaction cycle efficacy.

Figure 4c shows the mean bead fluorescence intensity for a completed ligation cycle compared to the negative control. Here, the negative control is an acceptor molecule with a 3’-hydroxyl where the click ligation is performed without azide addition. In the absence of the azide, the ligation step is expected to fail, and no fluorescence signal is expected for this negative control. The experimental results are consistent with this. In total, there are eight steps involving droplet encapsulation and decapsulation of the bead, which further demonstrates the robustness of the LAMPS for precise, multi-step biochemical processing.

## Discussion

We introduce LAMPS platform that implements picodroplet reactors with microbeads as solid supports to enable programmable, complex multi-step biochemical reactions at the picoliter scale. By engineering and leveraging OET for precise, nondestructive droplet and bead manipulation, we demonstrate sequential, controlled encapsulation and decapsulation of individual microbeads using the LAMPS platform. This establishes the ability of LAMPS to perform reliable reagent addition and removal across multiple reaction steps, supporting complex multi-step workflows. We validate these capabilities by performing an eight-step, template-independent oligonucleotide ligation cycle using modified click chemistry.

LAMPS overcomes the single-step limitations of current droplet microfluidic platforms by using light-induced virtual electrodes for dynamic, selective manipulation of droplets and beads with high spatial and temporal resolution. This enables on-demand control of reaction steps, concurrent processing of multiple bead-droplet pairs, and removes geometric restrictions associated with fixed electrodes. We also introduce a pulsed-voltage-based decapsulation mechanism that enables robust and repeatable bead decapsulation after each reaction step, a critical reagent removal capability for executing precise, multi-step solid-phase reactions.

Together, these features allow LAMPS to perform full-cycle, stepwise biochemical reactions with improved reproducibility and reliability.

Compared to conventional multi-step solid-phase reactors including column-based reactors, multi-well plates, and microarrays, which operate at microliter to milliliter scales, LAMPS offers a fundamentally different approach. Its droplet-based configuration reduces the minimum reaction volume by up to a million-fold, enabling reactions to proceed in picoliter-scale compartments. This is particularly valuable for applications such as assay development, prototyping, or low-volume synthesis, where only small amounts of product are required. In these cases, traditional platforms often require producing excess material to meet minimum volume constraints, resulting in unnecessary reagent consumption and waste. In addition to lowering the minimum viable reaction scale, LAMPS also achieves approximately a 25-fold decrease in the reagent-to-product ratio compared to standard benchtop reactions. This provides a more objective measure of reagent efficiency and remains advantageous independent of the product amount being produced, aligning with cost reduction and sustainability goals.

While LAMPS is not designed to match the throughput of bulk systems, its strengths lie in programmable control, precise manipulation of individual droplets, efficient reagent exchange, and parallel execution of reactions at the level of single beads. Unlike traditional platforms that rely on batch-processing, LAMPS enables site-specific control over each reaction, supporting diverse workflows, flexible timing, and condition-specific processing. This fine-grained control makes it well-suited for adaptive experimentation and integration with automated systems. By ensuring uniform reagent exposure, enabling effective washing, and supporting in situ fluorescence-based detection, LAMPS addresses longstanding challenges in solid-phase synthesis. Its compatibility with diverse reagents, demonstrated through the complete eight-step ligation cycle, suggests broader applicability to multi-step reactions beyond nucleic acid synthesis. In particular, its potential to combine synthesis, screening, and optimization within a single miniaturized device, together with compatibility with feedback-driven automation pipelines, positions LAMPS as a cost-effective alternative to liquid-handling robots in current life science laboratories. This makes it especially valuable when working with expensive, sensitive, or limited reagents, and opens avenues for applications in synthetic biology, molecular engineering, drug discovery, and personalized therapeutics.

## Methods

### Device design and fabrication

#### Step emulsification on-demand droplet generator

The droplet generators are fabricated using soft lithography techniques. To prepare the mask, a 10 μm-thick layer of SU-8 photoresist is spin-coated onto a bare silicon wafer, exposed to 365 nm UV light to define the four 50 μm-wide microfluidic channels, and post-exposure baked (PEB). The wafer is then coated with a 100 μm SU-8 layer, aligned to the alignment marks, and exposed to UV light to define the chamber (∼5 mm × 5 mm with filleted corners) and the oil inlet and outlet channels (600 μm wide). The sharp transitions between 10 μm-high channels and the 100 μm-high chamber facilitate on-demand droplet generation via Rayleigh-Plateau instability^21^. After another PEB, the wafer is developed in an SU-8 developer bath. Finally, the SU-8 mold is hard-baked. A 10:1 w/w mixture of PDMS prepolymer and curing agent is cast against the mold to fabricate the microfluidic component of the device.

#### Planar optoelectronic tweezers (OET)

The planar electrodes of OET are fabricated through bilayer lift-off process. A glass wafer is cleaned with piranha solution and primed with hexamethyldisilazane (HMDS). The wafer is then spin-coated with ∼250 nm of Lift-Off Layer 2000 (LOL 2000), baked at 170°C for 5 minutes, and coated with a 1 μm-thick layer of SPR3612 photoresist. The photoresist is then patterned (Hidelberg MLA 150, 120 mJ/cm²) to define the interdigitated electrode patterns (10 μm wide, 100 μm spacing). This configuration generates an electric field gradient sufficient for dielectrophoretic (DEP) trapping and manipulation of ∼50 μm-diameter reagent droplets in hexadecane. After developing, a 15 nm-thick chromium adhesion layer and a 200 nm gold layer are deposited by e-beam evaporation. The wafer is immersed in a Remover 1165 bath overnight and heated to 65°C for 30 minutes to lift-off excess material.

Before amorphous silicon deposition, the wafer is plasma cleaned for 1 min to remove organic contaminants (15 sccm O2 flow, 15 Pa pressure, 250 W RF power). A ∼1 μm-thick layer of hydrogenated amorphous silicon (a-Si:H) is deposited on the wafer using plasma enhanced chemical vapor deposition (PECVD) with 700 sccm SiH4 (5% in He), 1000 sccm He flow, 1500 mTorr pressure, 25 W RF power, at 300°C.

#### Device assembly and surface treatment

The PDMS piece containing the droplet generators and chamber structure is bonded to the planar OET substrate using oxygen plasma bonding (2 sccm O2 flow, 85 mTorr, 40 W RF power, 40 s). The bonded device is left under ambient conditions overnight to restore surface hydrophobicity, which is essential for reagent-in-oil droplet generation. Prior to experiments, the device is heated at 150-170°C to enhance the hydrophobicity of the a-Si surface. This treatment helps retain droplet integrity by preventing excessive spreading during trapping and manipulation.

### Experimental setup

The experimental setup, with the model numbers of the components labeled, is shown in Extended Data Fig. 1. The setup consists of optical, electrical and fluidics systems.

The *optical system* consists of micro-pattern projection and imaging subsystems. The micro- pattern projection subsystem uses a custom inverted microscope with a spatial light modulator (SLM) to project programmable light patterns onto the device photoconductor surface. The imaging subsystem enables both bright-field and fluorescence imaging.

The imaging subsystem is a modified upright epifluorescence microscope. To monitor experiments in bright field mode, transmitted light from the SLM through the device collected by a top objective lens and directed to a scientific CMOS (sCMOS) camera. For fluorescence imaging, a shutter blocks the light from the SLM, and sample illumination is provided through the top objective lens by an LED coupled with an excitation filter. Emission light is collected by the same lens, filtered through an emission filter, and captured by the camera.

The *electrical system* is composed of a waveform generator (i.e., signal generator), a radio frequency (RF) amplifier, and an oscilloscope to monitor voltage signals. To ensure sufficient voltage levels for effective trapping and manipulation, the signal from the waveform generator is amplified by the RF amplifier before being directed to the interdigitated electrodes of the device.

The *fluidics system* provides pressure pulses for the droplet generators and regulates the oil inlet fluidic port. A piezoelectric pressure controller, connected to a pressurized nitrogen line and a vacuum pump, delivers precise pressure to fluidic reservoirs, enabling controlled reagent flow through microfluidic tubing attached to the device. A total of four fluidic reservoirs and corresponding tubing (each connected to separate channels on the pressure controller) allow for independent control of reagent flow and droplet generation. For simplicity, only one of these reservoirs is illustrated in Extended Data Fig. 1.

### Binding of DNA acceptor to magnetic beads

Streptavidin-coated magnetic beads (EPRUI, MagSA-5, 5 μm in diameter, ∼6300 beads/μL) are used as solid supports. A 100 μL aliquot of beads is washed twice with 1 mL PBS. An acceptor solution is prepared by diluting 5 μL of 100 μM 5’-biotinylated DNA acceptor (sequences listed in Extended Data Table 1) with 95 μL of PBS (pH 7.4). The mixture of bead and acceptor is gently agitated using a tube rotator at 20 RPM for one hour to facilitate binding. Following incubation, the beads are washed multiple times with PBS and resuspended in 100 μL of PBS. The amount of DNA molecule bound per bead is estimated to be approximately 500 attomoles per bead.

### Suspending beads in hexadecane

A 5 μL aliquot of beads bound with DNA acceptor is placed on a magnetic stand, and the supernatant is carefully removed to isolate the beads. The Eppendorf tube with isolated beads is left open for 20 minutes to air dry, removing any residual PBS. Hexadecane containing 4% v/v Span 80 is then added as the suspension medium. The bead in hexadecane solution is vortexed to create a uniform colloidal solution.

### Droplet encapsulation and decapsulation of beads

Approximately 25 μL of each reagent is loaded into polyethylene microtubings (I.D. x O.D.: 0.38 mm x 1.09 mm) and connected directly to the corresponding reagent inlets on the device.

Individual droplets are generated at the sharp step transition between the shallow droplet generation channel and the deeper chamber by applying a pressure pulse of ∼20 mbar to the droplet generator. Once a droplet is generated near the droplet generator opening, a pair of rectangular light patterns are projected onto the device to seize the left and right sides of the droplet, aligning with the interdigitated electrode orientation.

To trap and manipulate the droplet in two dimensions, a sinusoidal AC voltage (9.3 V, 10 kHz) is applied to the interdigitated electrodes. The droplet is trapped in the gap between the rectangular light patterns and moves in tandem with the rectangle pair. The trapping force generated by the virtual electrodes activated by the light patterns is sufficient to overcome the droplet’s surface tension, allowing the bead to be fully encapsulated within the droplet. For decapsulation, the droplet is positioned at the central of a pair of physical electrodes. The voltage is then turned off, and the rectangular light patterns are moved slightly away from the droplet along the physical electrode pair. A pulse-modulated sinusoidal voltage (50% duty cycle, 2-second period), similar to amplitude shift keying (burst mode), is applied. During the active phase of the duty cycle, the sinusoidal signal has a frequency of 20 kHz with a magnitude of 40 V. The left and right sides of the droplet are abruptly pulled toward the rectangular light patterns without breaking up the droplet. During the reduced voltage period, the droplet retracts to its spherical shape, leaving the bead behind and completing the decapsulation process.

### Click ligation step and wash steps

Detailed reagent recipes are provided in Extended Data Table 2. The click ligation step starts with a bead bound with 3’-N3 DNA acceptors. The ligation process proceeds by generating two droplets: Click ligation (Part A) and Click ligation (Part B). Part A, which contains Atto 647- labeled donor molecules (sequence listed in Extended Data Table 1), is fluorescent, whereas Part B is non-fluorescent. Using two pairs of rectangular light patterns, these two droplets are maneuvered into proximity until they merge. The entire droplet becomes fluorescent immediately after merging, indicating thorough mixing of the components. After the bead is encapsulated by this merged droplet, it is incubated for 30 minutes before decapsulation. To enhance reagent exposure, the droplet is dragged around to stimulate internal fluid flow around the bead. This step is followed by three wash steps. In each wash step, the bead is encapsulated with a fresh PBS droplet (i.e., droplet from one wash step is not reused for the next wash step) for at least 30 seconds and then decapsulated. During each wash step, the droplet is moved back and forth by the light patterns to facilitate effective removal of unreacted materials. Fluorescence intensity of the bead decreases progressively over the first three wash steps, with no further reduction beyond the third wash. Therefore, the final fluorescence intensity of the bead is recorded after the third wash. For both the reaction and the negative control (using beads bound with 3’-OH DNA acceptors), the experiments are repeated five times. The mean values are shown in Fig. 4C.

### Click ligation synthesis cycle

The click ligation synthesis cycle starts with a bead bound with 3’-OH DNA acceptors, which is encapsulated in an enzymatic azide addition droplet (Extended Data Table 2) for 30 minutes, then decapsulated. This is followed by three PBS wash steps to remove unreacted components, resulting in 3’-N3 DNA product remain tethered to the bead. The wash steps are identical to the ones discussed in the previous section. The click ligation step then follows as previously described, with the final fluorescence intensity recorded after another three washes. Three data points are collected to calculate the mean value for the bar plot in Fig. 4C. While the demonstration on LAMPS focused on one full cycle, the iterative nature of the ligation synthesis cycle, which extends DNA primer molecules by defined sequences, indicates the method’s capability to produce extended oligonucleotides by simply repeating the steps already demonstrated.

### Benchtop reactions and polyacrylamide gel electrophoresis (PAGE) analysis

Click ligation reactions, modified from established protocols^40^, are initially tested in benchtop with bulk beads to validate benchtop performance before transitioning to on-chip reactions, following the same recipes outlined in Extended Data Table 2. Benchtop reactions are performed in 1.5 mL Eppendorf tubes, with bulk reagent handling instead of the droplet encapsulation and decapsulation of individual beads. To perform benchtop reactions, 5 µL of DNA acceptor-bound beads are placed on a magnetic stand, the supernatant is removed, and 50 µL of reagent (enzymatic azide addition or Click ligation A & B mixture) is added. The mixture is vortexed briefly and incubated for 30 minutes with periodic agitation. Beads are washed three times with 50 µL PBS and resuspended in 50 µL water. A 0.5 µL aliquot is used for fluorescence microscopy (Extended Data Figs. 2, 3).

For PAGE analysis, DNA is cleaved from beads by heating in 10 µL pure water at 80°C for 20 minutes. The supernatant is mixed with 10 µL 2x TBE-Urea buffer and heated to 80°C for 5 minutes. 8 µL of each sample is loaded onto a 15% TBE PAGE gel. Electrophoresis of the samples is performed at 100 V for 120 minutes. The gel is rinsed and imaged using the Cy5 channel on a gel imaging system, which can detect Atto 647 fluorescence. Bands representing full-length products are detected due to Atto 647 labeling of donor molecules (Extended Data Fig. 4).

### Fluorescence image analysis

Fluorescence images are captured in monochrome at 1920 × 1080 resolution and saved as 16-bit TIFF files to preserve detail. Example images of beads after benchtop reactions are shown in Extended Data Fig. 2. Images are processed using a custom code that applies adaptive thresholding to convert gray-scale images to binary masks, compensating for background variations by subdividing the image into blocks^45^. The resulting binary image, referred to as the mask, is shown in Extended Data Fig. 2b. Individual beads are identified using image segmentation and a single orthogonal hop connectivity criterion^46^. Each segmented region (sub- mask) corresponds to a detected bead. The segments are labeled on the mask in Extended Data Fig. 2c and on the fluorescence image in Extended Data Fig. 2d. These sub-masks are shown in Extended Data Fig. 3a. Multiplying each sub-mask with the original image extracts the pixels for the corresponding bead, as segmented images shown in Extended Data Fig. 3b. Overlapping beads are grouped into a single sub-mask, but size normalization ensures no errors.

The raw fluorescence intensity value of the 𝑖-th object, 𝐼_!,raw,#_, is calculated as:

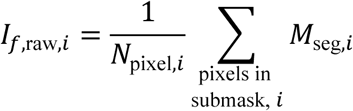

where 𝑖 = 1, 2, ⋯, 𝑁_$_, 𝑁_$_ is the total number of detected objects, 𝑁_pixel,#_ is the number of non-zero pixels in the 𝑖-th object, 𝑀_seg,#_ is the pixel value matrix of the 𝑖-th object, and each matrix element ranges from 0 to 65,535. This summing and averaging process automatically scales for bead size variations (e.g., a cluster of two beads).

The background intensity, 𝐼_&’_, is calculated by averaging from pixels outside detected objects. The corrected fluorescence intensity is then:

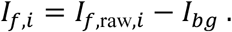

### Modeling and simulations

The modeling and simulation efforts include electromagnetic simulations to determine the field distribution near the virtual electrodes and resulting OET gradient force, and fluid flow simulations to track droplet motion under this force.

#### Electromagnetic simulations

The field distribution near the virtual electrodes depends on the geometry, excitation AC voltage, and material properties, with relevant parameters listed in Extended Data Table 3. The key material properties, permittivity and electrical conductivity, are sourced from the literature ^27,47–52^. Simulations are performed using the Electric Current interface (AC/DC module) of COMSOL Multiphysics, with electric potentials applied as Dirichlet boundary conditions at the physical electrodes and two rectangular virtual electrodes modeling the light patterns. The simulation setup is shown in Extended Data Fig. 5a, with the conductivity profiles of the system materials depicted in Extended Data Figs. 5b and 5c.

The electromagnetic field distributions obtained from the simulation (Fig. 2c) allow for calculation of the force on the droplet using the Maxwell Stress Tensor (MST) method ^53,54^. Force components along x, y and z directions (Extended Data Fig. 6) confirm that the droplet is attracted toward the virtual electrode gap, with sufficient gradient force generated near the electrodes to enable droplet manipulation.

#### Fluid flow simulations

To observe droplet motion under the gradient force, two-dimensional laminar two-phase flow simulations are performed using the phase-field method. The phase-field approach uses the variable “volume fraction” to represent the two fluids: values of 0 and 1 correspond to the droplet and the suspension medium, respectively, whereas intermediate values represent the boundary/transition region between the fluids. The fluid motions are governed by the Navier-Stokes equation in the laminar flow regime. Relevant material properties are listed in Extended Data Table 3. The interfacial surface tension between the droplet (PBS) and the medium (hexadecane) is taken to be 53.1 mN/m. All the material parameter values are taken from the literature^50,55–58^. Since PBS shares similar fluidic properties to water, water parameter values are used when PBS-specific data are unavailable.

In the two-phase flow, wetted-wall Navier-slip boundary conditions are set at the top and bottom surfaces of the fluid domain. The dielectrophoretic gradient force is coupled into the Multiphysics simulation as a volume force term defined in terms of the MST. The pressure field and the velocity field are initialized to zero. The droplet (radius 𝑟_()_ = 25 µm) starts at 𝑥 = −32 µm, 𝑧 = 35 µm (10 µm above the bottom surface), while a bead is positioned at 𝑥 = 0, 𝑧 = 5 µm. A time-dependent fluid-flow simulation is performed to track the position of the droplet as it moves under the gradient force (Fig. 2d).

## Data Availability

All data generated or analyzed during this study are included in this article (and its supplementary information files).

## Code Availability

The custom image processing code for fluorescence image analysis is available on GitHub at https://github.com/zaman13/16-bit-TIF-image-viewer.

## Acknowledgments

This research was funded by the National Institutes of Health (grant R01GM138716) and unrestricted research funding from Stanford University for Lambertus Hesselink. The devices were fabricated at the Stanford Nano Shared Facilities (SNSF)/Stanford Nanofabrication Facility (SNF), supported by the National Science Foundation under award ECCS-2026822, and at Stanford Microfluidics Foundry (SMF). Some of the illustrations were created in BioRender (https://BioRender.com). We thank Yao-Te Cheng, Jennifer Ortiz-Cárdenas, Thomas Eugene Carver, Sebastian Somolinos Cedeno, Drew Endy, Ludwig Galambos, Yasser Gidi, Peter Szijj, and Yue Zhang for helpful discussions.

## Author contributions

M.W. contributed to the design and fabrication of OET and droplet generator, the design and construction of the experimental setup, on-chip chemical reaction development, on-chip and benchtop experiments, data analysis, and light pattern programming code. M.A.Z. contributed to OET design and fabrication, the design and construction of the experimental setup, on-chip experiments, data analysis, light pattern programming code and fluorescence image processing code. M.A.J. contributed to the chemical reaction development and benchtop experiments. P.P. helped with droplet generator design. W.R. provided input for a-Si deposition. L.H., R.W.D., M.W., M.A.Z., and M.A.J. conceived the project. L.H. supervised the project. M.W. wrote the manuscript with inputs and edits from M.A.Z., L.H., M.A.J., and P.P.

## Materials & Correspondence

Correspondence and material requests should be addressed to Mo Wu.

## Supplementary information

Supplementary Information is available for this paper.

---

**Extended Data Fig. 1:**
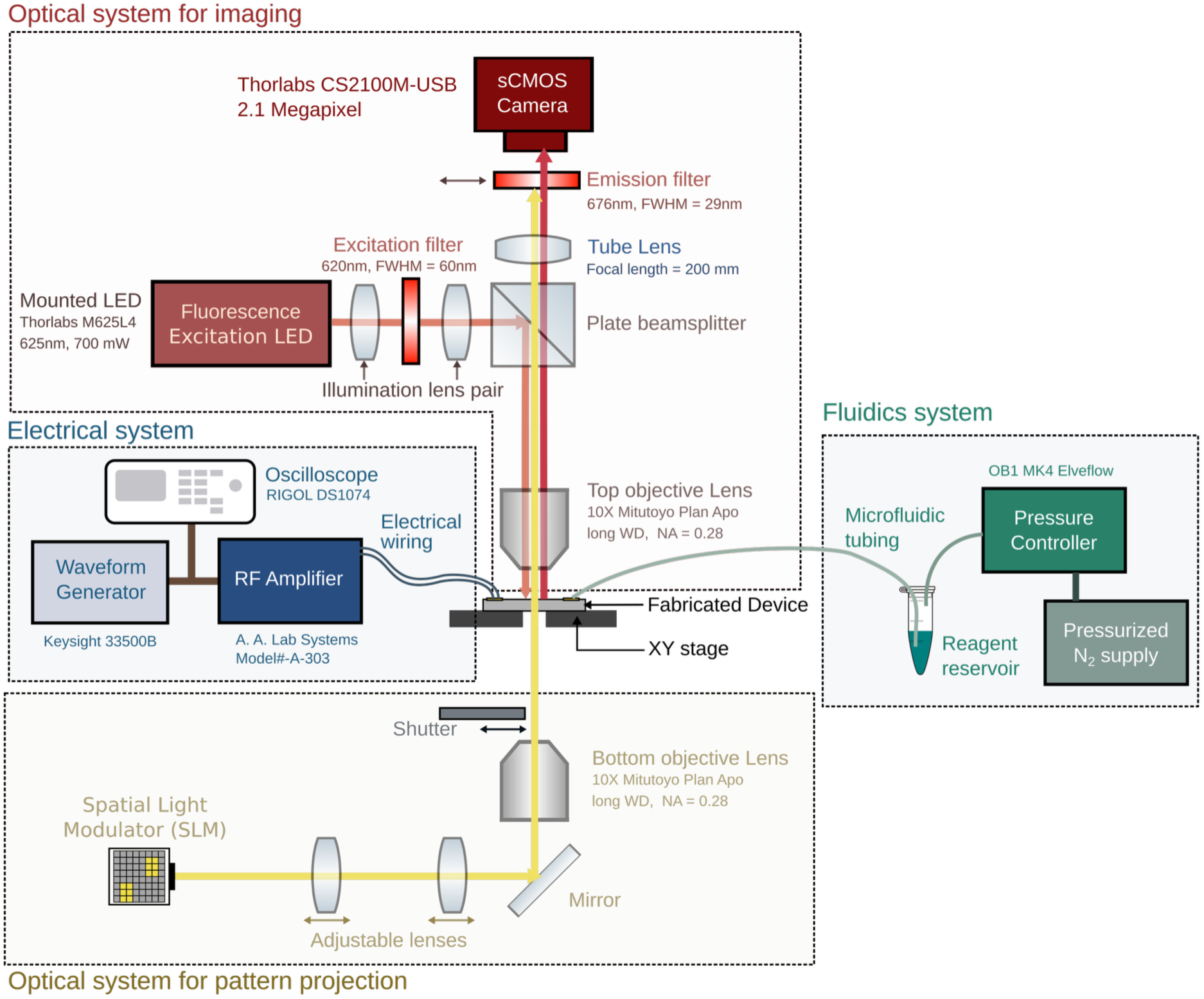
Schematic diagram of the experimental setup. Simplified optical paths are shown using colored arrows.

**Extended Data Fig. 2:**
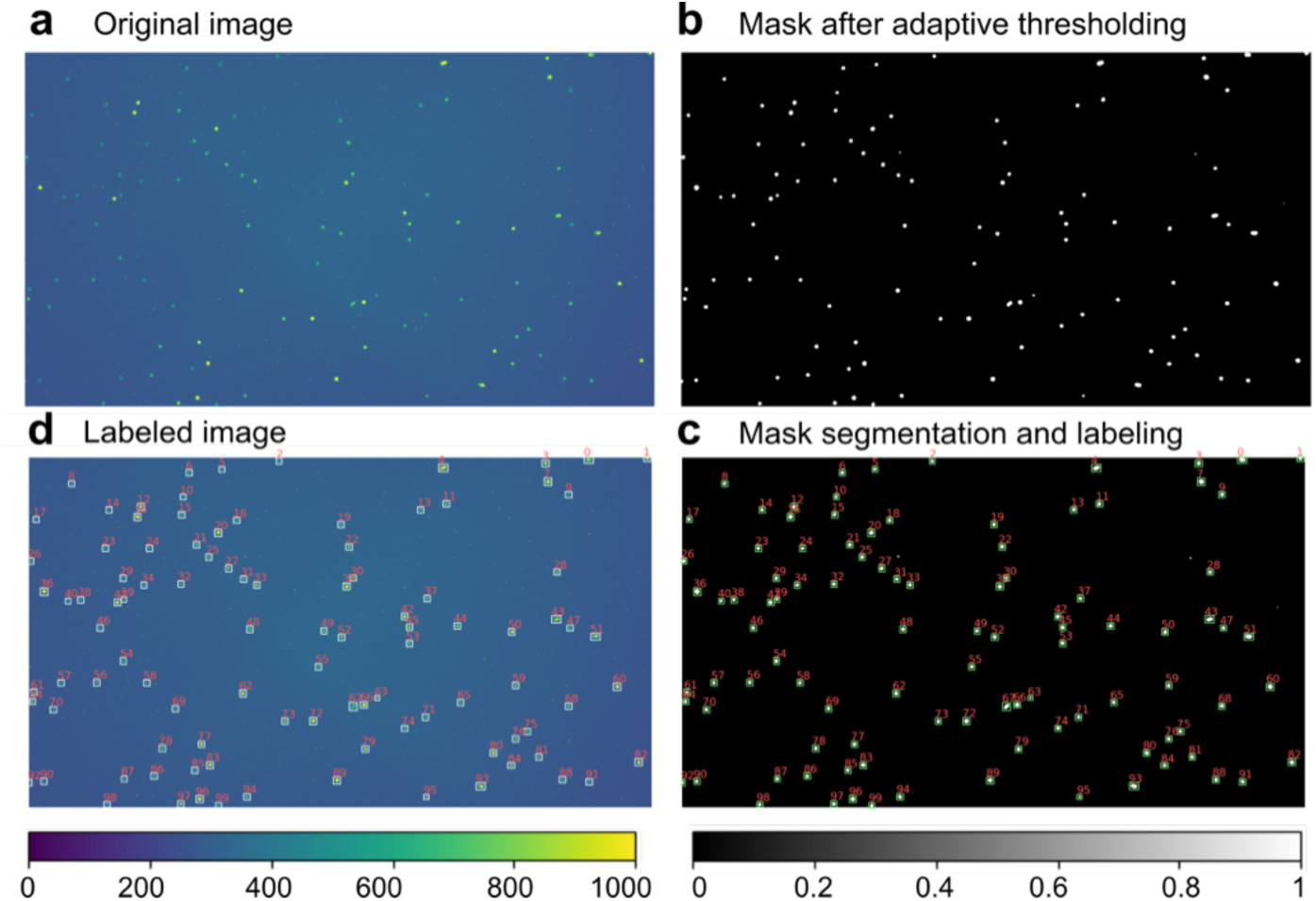
Fluorescence analysis. **a**, Original image. **b**, Binary image mask after adaptive thresholding. **c**, Labeled and segmented mask. **d**, Image with individual detected objects labeled.

**Extended Data Fig. 3:**
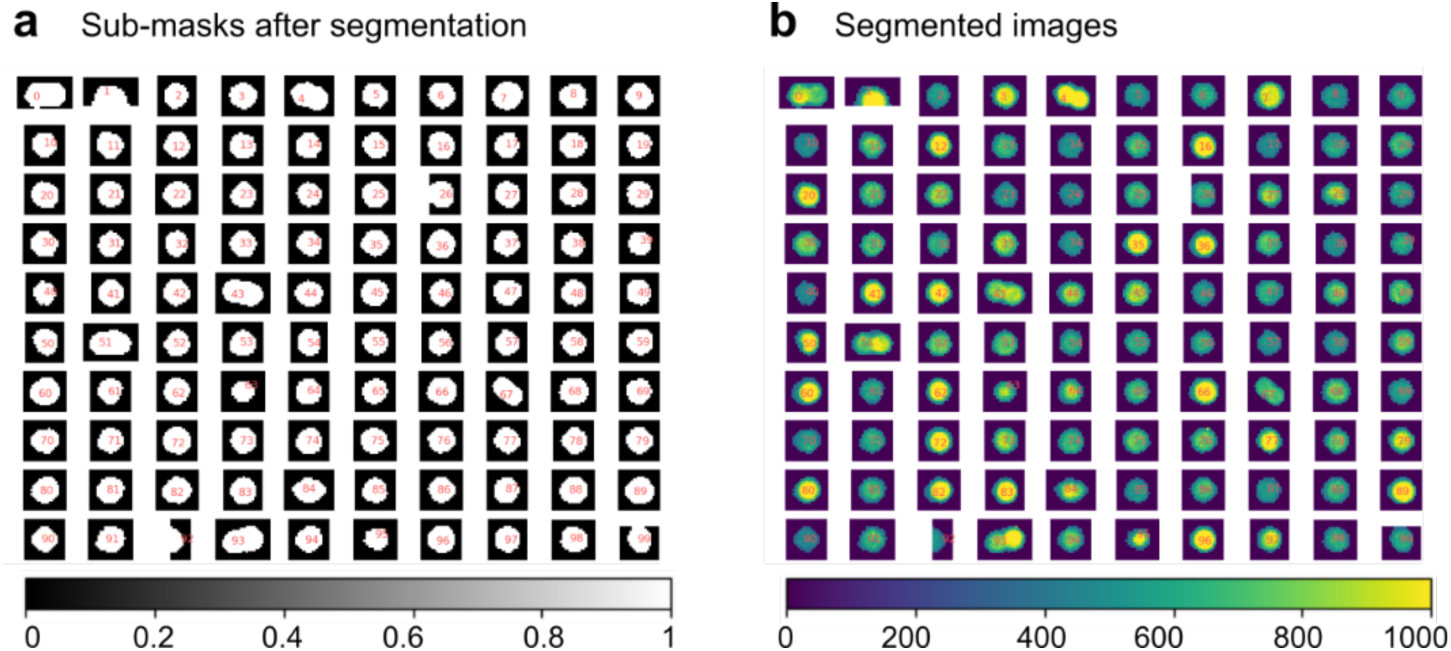
Sub-masks and segmented images. **a**, Binary sub-masks from segmentation. **b**, Image segments (sub-masks) corresponding to each detected object.

**Extended Data Fig. 4:**
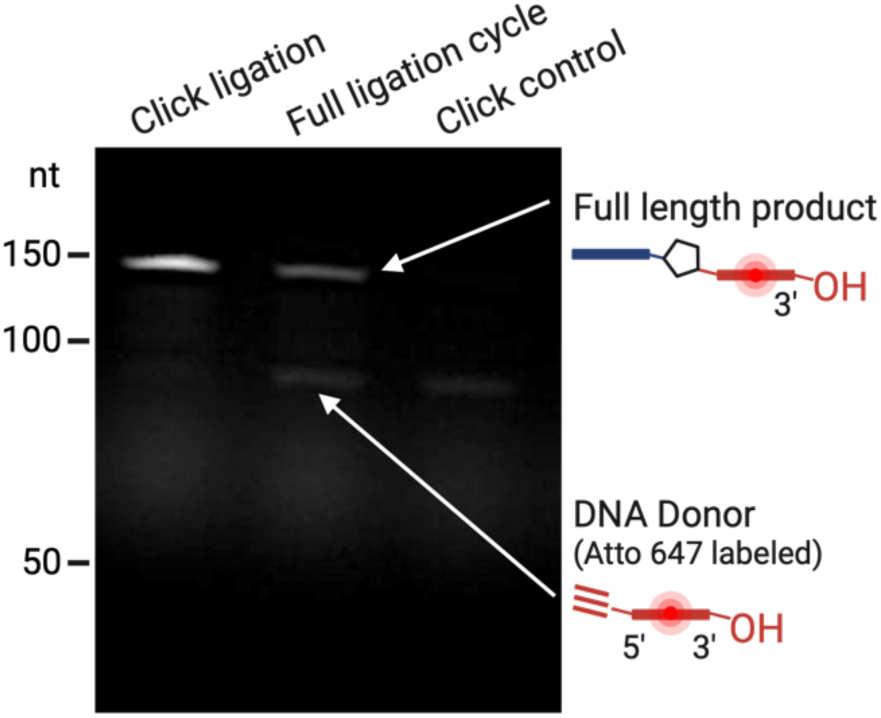
PAGE gel image in the Cy5 channel. It shows Atto 647-labeled products cleaved from beads after: (i) single click ligation step, (ii) full ligation cycle, and (iii) negative control.

**Extended Data Fig. 5:**
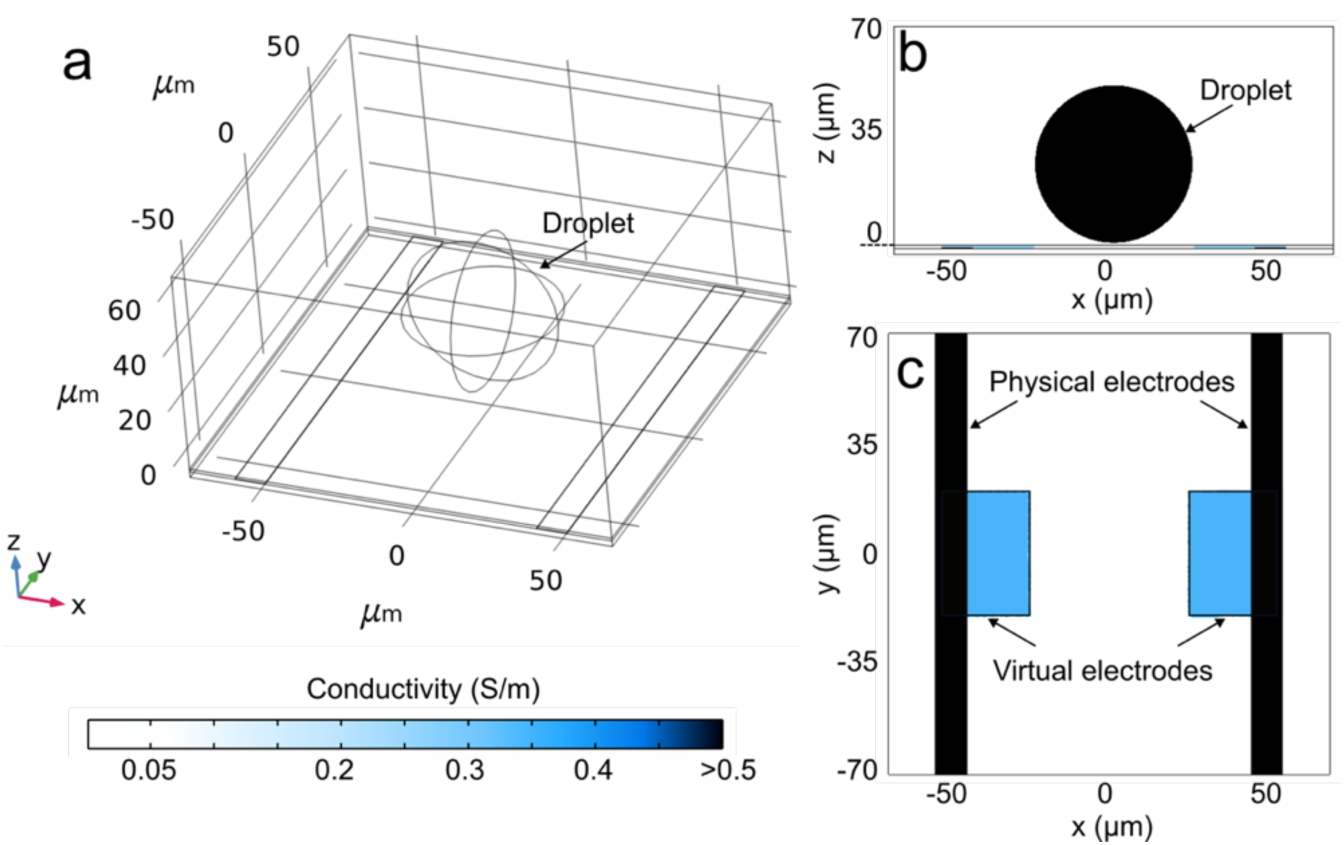
Geometry and conductivity values in the simulation system. 3D simulation geometry (**a**), with conductivity values of materials shown in the xz plane (**b**) and the xy plane (**c**). (**b**) and (**c**) share the same color bar. The virtual electrodes are not visible in the 3D geometry (**a**) as they are modeled as a conductivity function within the material properties rather than as a physical structure.

**Extended Data Fig. 6:**
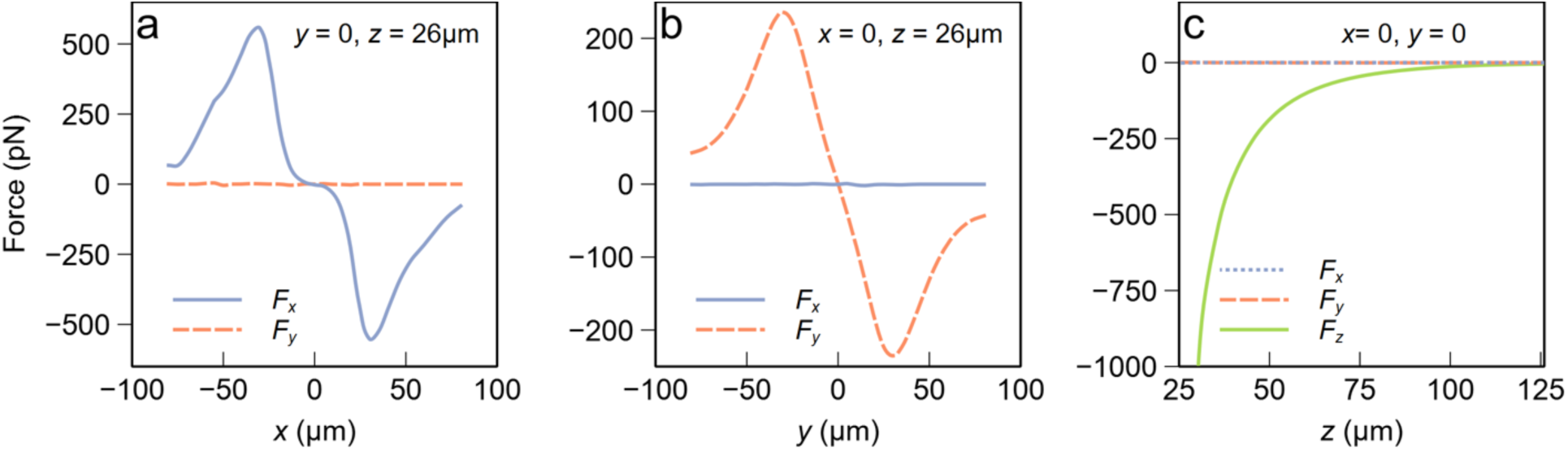
Force distribution near the virtual electrodes. The droplet position is swept in the x (**a**), y (**b**), and z (**c**) directions to generate the plots, with the remaining coordinates held constant at the values indicated on each plot. Fx, Fy, and Fz represent the x, y, and z components of the force.

**Extended Data Table 1.**
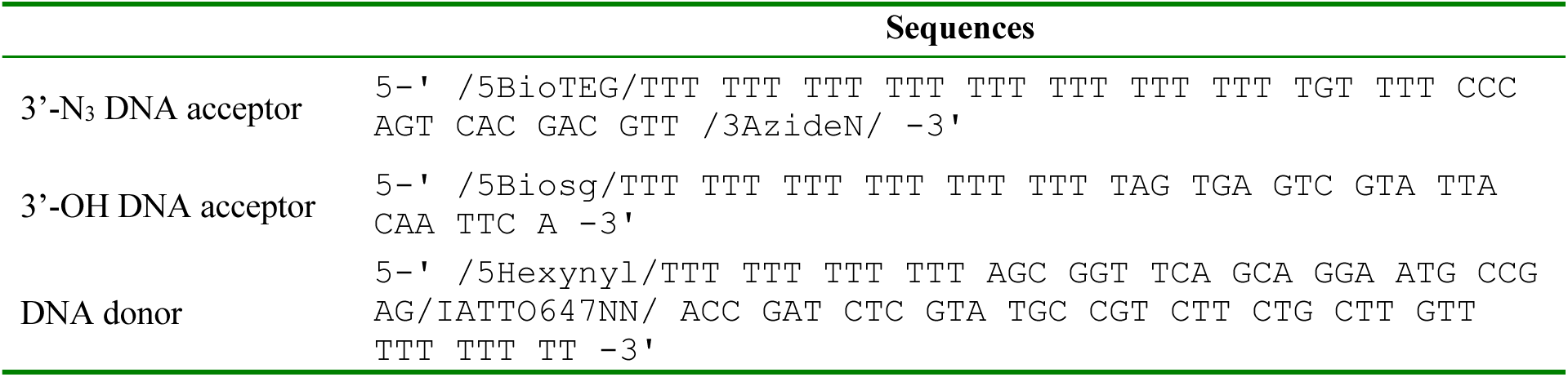
Sequences of DNA acceptor and donor strands used in click ligation reactions.

**Extended Data Table 2.** Click ligation synthesis cycle recipes.

| **Reagent name** | **Volume (μL)** |
| --- | --- |
| **Step 1: Enzymatic azide addition** |  |
| water | 31 |
| Terminal Transferase Reaction Buffer (10x, New England Biolabs) | 5 |
| CoCl<sub>2</sub> for TdT (2.5 mM, New England Biolabs) | 5 |
| Zidovudine Triphosphate (AzTTP, 10 mM, Jena Bioscience) | 3 |
| Terminal Transferase (TdT, 20 U/μL, New England Biolabs) | 6 |
| **Step 5: Click ligation (Part A)** |  |
| *tert*-Butanol (t-BuOH) | 19 |
| water | 1 |
| donor-Atto 647 (100 μM, IDT) | 5 |
| **Step 5: Click ligation (Part B)** |  |
| *tert*-Butanol (t-BuOH) | 19 |
| water | 4 |
| Cu(I) catalyst solution | 2 |
| **Steps 2 to 4 and 6 to 8: Washing steps** |  |
| PBS (pH 7.4) | 50 |
| **Cu(I) catalyst solution** |  |
| 25mM CuSO<sub>4</sub> | 5 |
| 125mM THPTA | 5 |
| 500mM NaAsc | 5 |
| water | 5 |

**Extended Data Table 3.**
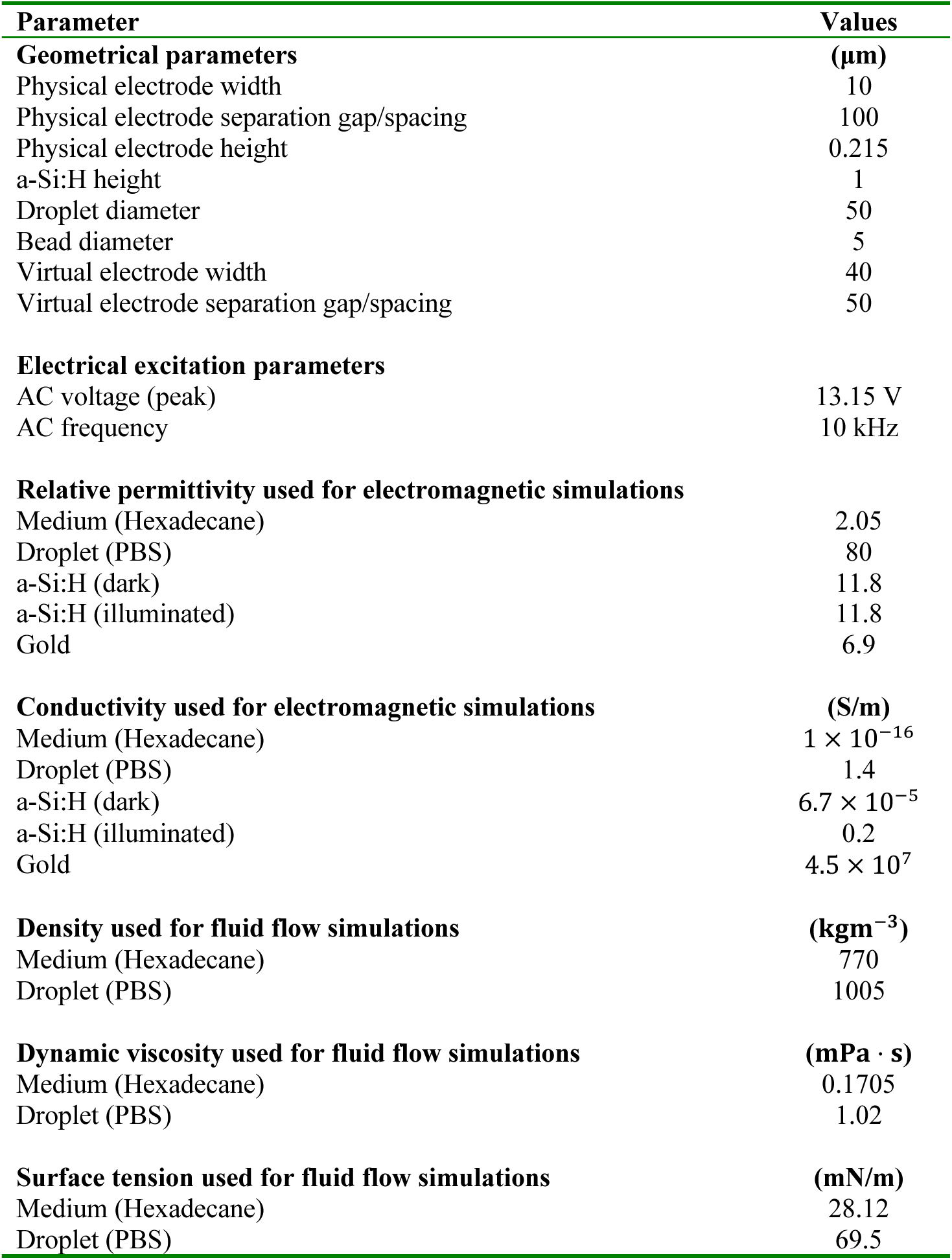
Parameters used in simulations.

## References

1 Shang, L., Cheng, Y. & Zhao, Y. Emerging Droplet Microfluidics. Chemical Reviews 117, 7964–8040 (2017). 10.1021/acs.chemrev.6b00848

2 Kaminski, T. S. & Garstecki, P. Controlled droplet microfluidic systems for multistep chemical and biological assays. Chemical Society Reviews 46, 6210–6226 (2017). 10.1039/C5CS00717H

3 Gach, P. C., Iwai, K., Kim, P. W., Hillson, N. J. & Singh, A. K. Droplet microfluidics for synthetic biology. Lab on a Chip 17, 3388–3400 (2017). 10.1039/C7LC00576H

4 Duncombe, T. A., Tentori, A. M. & Herr, A. E. Microfluidics: reframing biological enquiry. Nature Reviews Molecular Cell Biology 16, 554–567 (2015). 10.1038/nrm4041

5 Gines, G. et al. Functional analysis of single enzymes combining programmable molecular circuits with droplet-based microfluidics. Nature Nanotechnology 19, 800–809 (2024). 10.1038/s41565-024-01617-1

6 Macosko, Evan Z. et al. Highly Parallel Genome-wide Expression Profiling of Individual Cells Using Nanoliter Droplets. Cell 161, 1202–1214 (2015). 10.1016/j.cell.2015.05.002

7 Shembekar, N., Chaipan, C., Utharala, R. & Merten, C. A. Droplet-based microfluidics in drug discovery, transcriptomics and high-throughput molecular genetics. Lab on a Chip 16, 1314–1331 (2016). 10.1039/C6LC00249H

8 Agresti, J. J. et al. Ultrahigh-throughput screening in drop-based microfluidics for directed evolution. Proceedings of the National Academy of Sciences 107, 4004–4009 (2010). doi:10.1073/pnas.0910781107

9 Ma, Y., Zhang, Z., Jia, B. & Yuan, Y. Automated high-throughput DNA synthesis and assembly. Heliyon 10, e26967 (2024). 10.1016/j.heliyon.2024.e26967

10 Pon, R. T. Solid-Phase Supports for Oligonucleotide Synthesis. Current Protocols in Nucleic Acid Chemistry 00, 3.1.1–3.1.28 (2000). 10.1002/0471142700.nc0301s00

11 Cheng, J. Y., Chen, H. H., Kao, Y. S., Kao, W. C. & Peck, K. High throughput parallel synthesis of oligonucleotides with 1536 channel synthesizer. Nucleic Acids Res 30, e93 (2002). 10.1093/nar/gnf092

12 Coin, I., Beyermann, M. & Bienert, M. Solid-phase peptide synthesis: from standard procedures to the synthesis of difficult sequences. Nature Protocols 2, 3247–3256 (2007). 10.1038/nprot.2007.454

13 Afting, C. et al. DNA microbeads for spatio-temporally controlled morphogen release within organoids. Nature Nanotechnology 19, 1849–1857 (2024). 10.1038/s41565-024-01779-y

14 Wolfrum, C., Josten, A. & Götz, P. Optimization and scale-up of oligonucleotide synthesis in packed bed reactors using computational fluid dynamics modeling. Biotechnology Progress 30, 1048–1056 (2014). 10.1002/btpr.1966

15 Sletten, E. T., Nuño, M., Guthrie, D. & Seeberger, P. H. Real-time monitoring of solid-phase peptide synthesis using a variable bed flow reactor. Chemical Communications 55, 14598–14601 (2019). 10.1039/C9CC08421E

16 Padhy, P. et al. Dielectrophoretic bead-droplet reactor for solid-phase synthesis. Nature Communications 15, 6159 (2024). 10.1038/s41467-024-49284-z

17 Matuła, K., Rivello, F. & Huck, W. T. S. Single-Cell Analysis Using Droplet Microfluidics. Advanced Biosystems 4, 1900188 (2020). 10.1002/adbi.201900188

18 Witters, D., Knez, K., Ceyssens, F., Puers, R. & Lammertyn, J. Digital microfluidics-enabled single-molecule detection by printing and sealing single magnetic beads in femtoliter droplets. Lab on a Chip 13, 2047–2054 (2013). 10.1039/C3LC50119A

19 Shim, J.-u., et al. Ultrarapid Generation of Femtoliter Microfluidic Droplets for Single-Molecule-Counting Immunoassays. ACS Nano 7, 5955–5964 (2013). 10.1021/nn401661d

20 Dressman, D., Yan, H., Traverso, G., Kinzler, K. W. & Vogelstein, B. Transforming single DNA molecules into fluorescent magnetic particles for detection and enumeration of genetic variations. Proceedings of the National Academy of Sciences 100, 8817–8822 (2003). doi:10.1073/pnas.1133470100

21 Dangla, R., Fradet, E., Lopez, Y. & Baroud, C. N. The physical mechanisms of step emulsification. Journal of Physics D: Applied Physics 46, 114003 (2013). 10.1088/0022-3727/46/11/114003

22 Mittal, N., Cohen, C., Bibette, J. & Bremond, N. Dynamics of step-emulsification: From a single to a collection of emulsion droplet generators. Physics of Fluids 26 (2014). 10.1063/1.4892949

23 Zhu, P. & Wang, L. Passive and active droplet generation with microfluidics: a review. Lab on a Chip 17, 34–75 (2017). 10.1039/C6LC01018K

24 Park, S. et al. Floating electrode optoelectronic tweezers: Light-driven dielectrophoretic droplet manipulation in electrically insulating oil medium. Appl Phys Lett 92, 151101–1511013 (2008). 10.1063/1.2906362

25 Chiou, P. Y., Ohta, A. T. & Wu, M. C. Massively parallel manipulation of single cells and microparticles using optical images. Nature 436, 370–372 (2005). 10.1038/nature03831

26 Zhang, S. et al. Optoelectronic tweezers: a versatile toolbox for nano-/micro-manipulation. Chemical Society Reviews 51, 9203–9242 (2022). 10.1039/D2CS00359G

27 Zaman, M. A., Wu, M., Ren, W. & Hesselink, L. Impedance matching in optically induced dielectrophoresis: Effect of medium conductivity on trapping force. Applied Physics Letters 125 (2024). 10.1063/5.0223354

28 Thio, S. K., Bae, S. W. & Park, S.-Y. Lab on a smartphone (LOS): A smartphone-integrated, plasmonic-enhanced optoelectrowetting (OEW) platform for on-chip water quality monitoring through LAMP assays. Sensors and Actuators B: Chemical 358, 131543 (2022). 10.1016/j.snb.2022.131543

29 Thio, S. K. & Park, S.-Y. A review of optoelectrowetting (OEW): from fundamentals to lab-on-a- smartphone (LOS) applications to environmental sensors. Lab on a Chip 22, 3987–4006 (2022). 10.1039/D2LC00372D

30 Jones, T. B. On the Relationship of Dielectrophoresis and Electrowetting. Langmuir 18, 4437–4443 (2002). 10.1021/la025616b

31 Jones, T. B., Gunji, M., Washizu, M. & Feldman, M. J. Dielectrophoretic liquid actuation and nanodroplet formation. Journal of Applied Physics 89, 1441–1448 (2001). 10.1063/1.1332799

32 Lee, S. J., Hong, J., Kang, K. H., Kang, I. S. & Lee, S. J. Electrowetting-induced droplet detachment from hydrophobic surfaces. Langmuir 30, 1805–1811 (2014). 10.1021/la404344y

33 El-Sagheer, A. H. & Brown, T. Click Nucleic Acid Ligation: Applications in Biology and Nanotechnology. Accounts of Chemical Research 45, 1258–1267 (2012). 10.1021/ar200321n

34 Kollaschinski, M. et al. Efficient DNA Click Reaction Replaces Enzymatic Ligation. Bioconjugate Chemistry 31, 507–512 (2020). 10.1021/acs.bioconjchem.9b00805

35 Fantoni, N. Z., El-Sagheer, A. H. & Brown, T. A Hitchhiker’s Guide to Click-Chemistry with Nucleic Acids. Chemical Reviews 121, 7122–7154 (2021). 10.1021/acs.chemrev.0c00928

36 Presolski, S. I., Hong, V. P. & Finn, M. G. Copper-Catalyzed Azide-Alkyne Click Chemistry for Bioconjugation. Curr Protoc Chem Biol 3, 153–162 (2011). 10.1002/9780470559277.ch110148

37. Kolb, H. C., Finn, M. G. & Sharpless, K. B. Click Chemistry: Diverse Chemical Function from a Few Good Reactions. Angewandte Chemie International Edition 40, 2004-2021 (2001). 10.1002/1521-3773(20010601)40:11<2004::AID-ANIE2004>3.0.CO;2-5

38 Qiu, J., El-Sagheer, A. H. & Brown, T. Solid phase click ligation for the synthesis of very long oligonucleotides. Chemical Communications 49, 6959–6961 (2013). 10.1039/C3CC42451K

39 Isobe, H., Fujino, T., Yamazaki, N., Guillot-Nieckowski, M. & Nakamura, E. Triazole-Linked Analogue of Deoxyribonucleic Acid (TLDNA): Design, Synthesis, and Double-Strand Formation with Natural DNA. Organic Letters 10, 3729–3732 (2008). 10.1021/ol801230k

40 Miura, F. et al. Triazole linking for preparation of a next-generation sequencing library from single-stranded DNA. Nucleic Acids Res 46, e95 (2018). 10.1093/nar/gky452

41 Neumann, S., Biewend, M., Rana, S. & Binder, W. H. The CuAAC: Principles, Homogeneous and Heterogeneous Catalysts, and Novel Developments and Applications. Macromol Rapid Commun 41, e1900359 (2020). 10.1002/marc.201900359

42 Chen, X., Khairallah, G. N., O’Hair, R. A. J. & Williams, S. J. Fixed-charge labels for simplified reaction analysis: 5-hydroxy-1,2,3-triazoles as byproducts of a copper(I)-catalyzed click reaction. Tetrahedron Letters 52, 2750-2753 (2011). 10.1016/j.tetlet.2011.03.094

43 Palluk, S. et al. De novo DNA synthesis using polymerase-nucleotide conjugates. Nature Biotechnology 36, 645–650 (2018). 10.1038/nbt.4173

44 Jensen, M. A. & Davis, R. W. Template-Independent Enzymatic Oligonucleotide Synthesis (TiEOS): Its History, Prospects, and Challenges. Biochemistry 57, 1821–1832 (2018). 10.1021/acs.biochem.7b00937

45 Pai, Y.-T., Chang, Y.-F. & Ruan, S.-J. Adaptive thresholding algorithm: Efficient computation technique based on intelligent block detection for degraded document images. Pattern Recognition 43, 3177–3187 (2010). 10.1016/j.patcog.2010.03.014

46 Braga-Neto, U. & Goutsias, J. A Theoretical Tour of Connectivity in Image Processing and Analysis. Journal of Mathematical Imaging and Vision 19, 5–31 (2003). 10.1023/A:1024476403183

47 Salmanzadeh, A., Shafiee, H., Davalos, R. V. & Stremler, M. A. Microfluidic mixing using contactless dielectrophoresis. ELECTROPHORESIS 32, 2569–2578 (2011). 10.1002/elps.201100171

48 Schropp, R. E. I. Improved material properties of amorphous silicon from silane by fluorine implantation: Application to thin-film transistors. Journal of Applied Physics 65, 3706–3711 (1989). 10.1063/1.342598

49 Zhao, Y., Hu, S. & Wang, Q. Simulation and analysis of particle trajectory caused by the optical-induced dielectrophoresis force. Microfluidics and Nanofluidics 16, 533–540 (2014). 10.1007/s10404-013-1246-1

50 Fan, S.-K., Hsieh, T.-H. & Lin, D.-Y. General digital microfluidic platform manipulating dielectric and conductive droplets by dielectrophoresis and electrowetting. Lab on a Chip 9, 1236–1242 (2009). 10.1039/B816535A

51 Pei Yu, C., Wilson, W., Liao, J. C. & Wu, M. C. in *17th IEEE International Conference on Micro Electro Mechanical Systems. Maastricht MEMS 2004 Technical Digest.* 21-24.

52 Hsieh, T. H. & Fan, S. K. in 2008 *IEEE 21st International Conference on Micro Electro Mechanical Systems*. 641–644.

53 Zhang, S. et al. Escape from an Optoelectronic Tweezer Trap: experimental results and simulations. Opt. Express 26, 5300–5309 (2018). 10.1364/OE.26.005300

54 Zaman, M. A., Padhy, P., Cheng, Y.-T., Galambos, L. & Hesselink, L. Optoelectronic tweezers with a non-uniform background field. Applied Physics Letters 117, 171102 (2020). 10.1063/5.0020446

55 Haynes, W. M. CRC Handbook of Chemistry and Physics (97th ed.). (CRC Press, 2016).

56 Brown, P. H., Balbo, A., Zhao, H., Ebel, C. & Schuck, P. Density Contrast Sedimentation Velocity for the Determination of Protein Partial-Specific Volumes. PLOS ONE 6, e26221 (2011). 10.1371/journal.pone.0026221

57 Palasz, A. T., Thundathil, J., Verrall, R. E. & Mapletoft, R. J. The effect of macromolecular supplementation on the surface tension of TCM-199 and the utilization of growth factors by bovine oocytes and embryos in culture. Animal Reproduction Science 58, 229–240 (2000). 10.1016/S0378-4320(99)00090-1

58 Cortés-Estrada, A. H., Ibarra-Bracamontes, L. A., Aguilar-Corona, A., Viramontes-Gamboa, G. & Carbajal-De la Torre, G. in Experimental and Computational Fluid Mechanics (eds Jaime Klapp & Abraham Medina) 219-226 (Springer International Publishing, 2014).

